# t-Darpp is an elongated monomer that binds to calcium and is phosphorylated by cyclin-dependent kinases 1 and 5

**DOI:** 10.1101/152272

**Authors:** Jamil Momand, Patrycja Magdziarz, You Feng, Dianlu Jiang, Elizabeth Parga, Arianna Celis, Erin Denny, Xiaoying Wang, Martin L. Phillips, Estuardo Monterroso, Susan E. Kane, Feimeng Zhou

## Abstract

t-Darpp is a protein encoded by the PPP1R1B gene and is expressed in breast, colon, esophageal, gastric, and prostate cancers, as well as in normal adult brain striatal cells. Overexpression of t-Darpp in cultured cells leads to increased protein kinase A activity and increased phosphorylation of AKT (protein kinase B). In HER2+ breast cancer cells t-Darpp confers resistance to the chemotherapeutic agent trastuzumab. To shed light on t-Darpp function, we studied its secondary structure, oligomerization status, metal-binding properties, and phosphorylation by cyclin dependent kinases 1 and 5. t-Darpp exhibits 12% alpha helix, 29% beta strand, 24% beta turn and 35% random coil structures. t-Darpp binds to calcium, but not to other metals commonly found in biological systems. The T39 site, critical for t-Darpp activation of the AKT signaling pathway, is a substrate for phosphorylation by cyclin-dependent kinase 1 (CDK1) and cyclin-dependent kinase 5 (CDK5). Gel filtration chromatography, sedimentation equilibrium analysis, blue native gel electrophoresis, and glutaraldehyde-mediated crosslinking experiments demonstrate that the majority of t-Darpp exists as a monomer, but forms low levels (< 3%) of hetero-oligomers with its longer isoform Darpp-32. t-Darpp has a large Stokes radius of 4.4 nm relative to its mass of 19 kDa, indicating that it has an elongated structure.

## INTRODUCTION

t-Darpp (truncated isoform of Dopamine- and cAMP-Regulated Phosphoprotein) is expressed in adult brain striatal cells and in many types of cancers including those of breast, colon, esophageal, gastric and prostrate origin [1-5]. Upon overexpression of t-Darpp in cultured cells protein kinase A (PKA) becomes active and the AKT cell survival pathway is strengthened [1, 6-8]. Much work has been focused on t-Darpp’s role in cancers. In breast cancer, t-Darpp was observed to be overexpressed in some breast cancers [1]. One type of breast cancer, characterized by high levels of human epidermal growth factor receptor (HER2/neu/ERBB2), is sensitive to an antibody drug called trastuzumab (Herceptin). Overexpression of t-Darpp in HER2+ breast cancer cells causes these cells to become resistant to the anti-proliferative effects of trastuzumab [1, 2, 6, 8].

An isoform of t-Darpp, named Darpp-32, has an additional 36 amino acids on its N-terminus and is translated from a transcript produced from an alternative promoter that codes for an upstream transcription start site [9]. Darpp-32 is expressed in the brain, specifically in regions enriched in dopaminoceptive neurons [10, 11]. Depending on its phosphorylation state, Darpp-32 inhibits protein phosphatase 1 (PP-1) and PKA [12, 13]. In some studies this longer isoform has been shown to inhibit cell growth and reverse t-Darpp’s ability to confer trastuzumab resistance and PKA activation [8]. However, other studies have shown that Darpp-32 is capable of increasing cell growth [14] and confers trastuzumab resistance [2]. In this study, we report the first description of the structure properties of t-Darpp. t-Darpp is an elongated monomer that binds to calcium and is a substrate for phosphorylation by CDK1 and CDK5.

## RESULTS

### Oligomerization analysis

To assess the oligomerization properties of t-Darpp, recombinant human t-Darpp protein was analyzed by blue native gel electrophoresis. In this procedure, proteins were not treated with denaturing and reducing agents. Figure 1A shows that the majority of t-Darpp (calculated mass 19.4 kDa based on sequence) runs slightly smaller than the 20 kDa marker, as expected. The staining pattern of the gel shows very small amounts of t-Darpp with mass >20 kDa; one form runs between 20 kDa and 66 kDa and several forms run between ~80 kDa and 150 kDa. Bovine serum albumin (BSA) was used as a comparison. BSA is a 66.5 kDa protein with little propensity to form oligomers. As expected, we find that the majority of BSA runs close to the 66 kDa protein standard (Figure 1A). Also observed are two higher mass forms of BSA — one at ~130 kDa and one at higher mass, produced by infrequently formed dimers and trimers, respectively [15].

**Figure 1.**
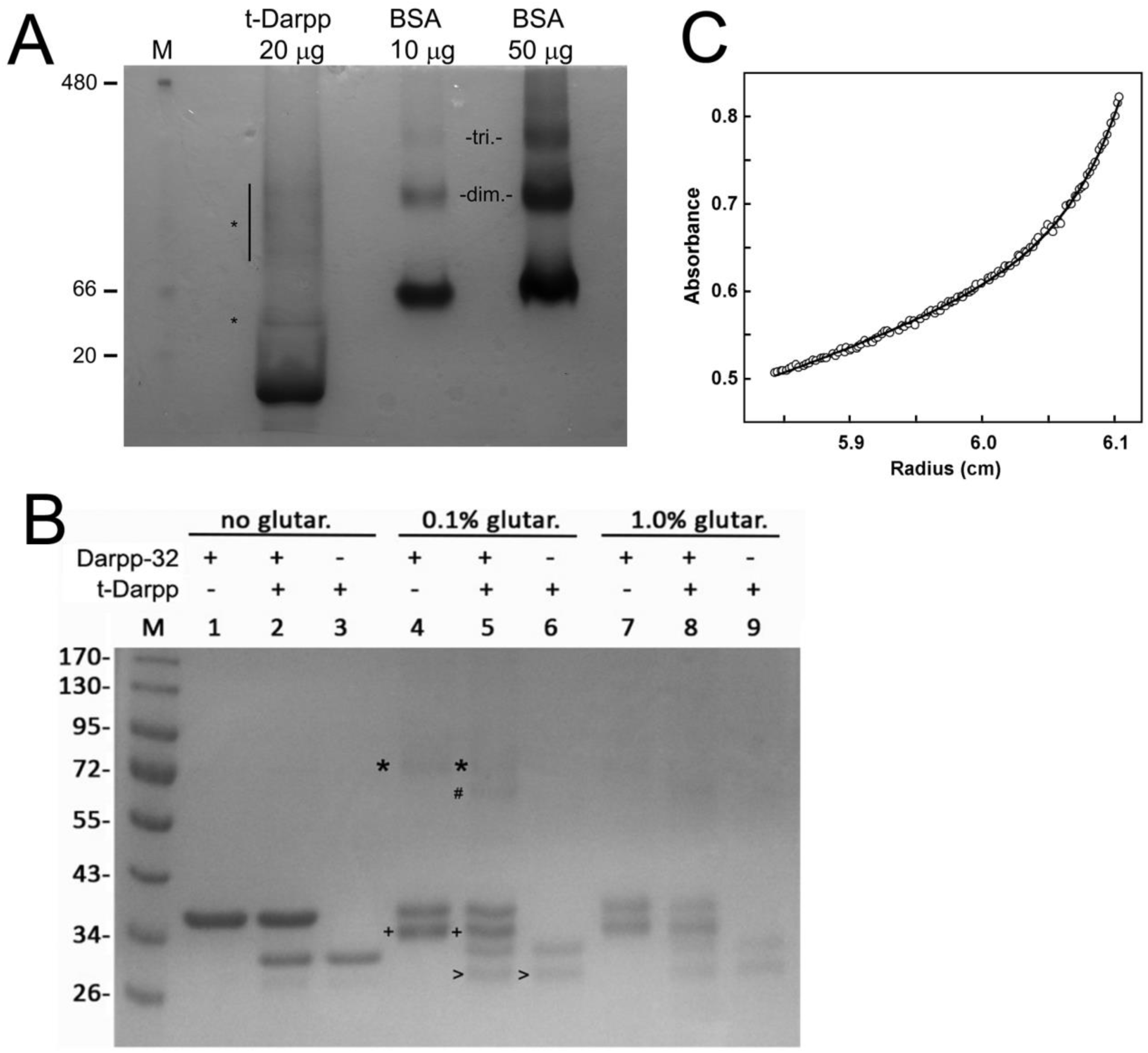
Oligomer analysis of t-Darpp and Darpp-32. A, Blue native gel analysis of t-Darpp.First lane, molecular weight markers (in kDa). Second lane, t-Darpp. Asterisks indicate higher mass forms of t-Darpp. Third lane, BSA (20 μg) and fourth lane, BSA (50 μg). The abbreviations “dim.” and “tri.” refer to dimer and trimer respectively. B, Glutaraldehyde treatment of t-Darpp, Darpp-32, and a mixture of t-Darpp and Darpp-32. Lanes 1-3, no treatment with glutaraldehyde; lanes 4-6, treatment with 0.1% glutaraldehyde; lanes 7-9, treatment with 1% glutaraldehyde. Asterisk denotes Darpp-32 homo-dimers; # denotes t-Darpp/Darpp-32 heterodimer; + denotes intramolecular crosslink in Darpp-32; > denotes intramolecular crosslink in t-Darpp. C, Sedimentation equilibrium of t-Darpp. Absorbance at 280 nm vs. radius (cm) plotted and line generated with a function that shows 96% monomer and 4% oligomer.

It has been reported that t-Darpp has some positive growth effects, likely by activating the PKA and/or the AKT pathways [1, 8]. One study has shown that its longer isoform, Darpp-32, has some growth inhibitory effects [8]. To explore the possibility that t-Darpp may inhibit Darpp-32 activity through direct protein-protein interaction they were tested for heterooligomeric complex formation. An equimolar mixture of recombinant t-Darpp and recombinant Darpp-32 was treated with glutaraldehyde, a primary amine crosslinking agent, and analyzed by SDS-PAGE (Figure 1B).

Without glutaraldehyde, Darpp-32 has an apparent mass of 36 kDa on SDS-PAGE and t-Darpp has an apparent mass of 30 kDa. These are significantly higher than their actual masses and are likely to be due to the high frequency of acidic residues in these proteins. Glutaraldehyde treatment of recombinant Darpp-32 produced a low level of high-mass species of approximately 72 kDa. Glutaradehyde treatment of the Darpp-32/t-Darpp mixture produced a low level of two high-mass species -- one approximately 72 kDa and the other approximately 68 kDa. Glutaraldehyde treatment of t-Darpp alone did not produce detectable levels of high-mass species. Thus, the 68 kDa species appears to be specific to the Darpp-32/t-Darpp combination and its size is consistent with such a hetero-dimer composition. Quantification of band intensities indicates that only about 3% of the Darpp-32 can form a cross-linked homo-oligomer or a hetero-oligomer with t-Darpp. No detection of t-Darpp homo-oligomers was observed, largely in agreement with blue native gel analysis.

Although cross-linking showed little dimerization, Figure 1B shows that glutaraldehyde treatment of Darpp-32 alone produced a significant level of a protein species with an apparent mass (34 kDa) lower than the untreated Darpp-32 (36 kDa). Similarly, treatment of t-Darpp with the cross-linker produced a significant level of a species with an apparent mass (27 kDa) lower than the untreated t-Darpp (30 kDa). The presence of these apparent lower mass proteins suggests that both t-Darpp and Darpp32 form intramolecular cross-links with glutaraldehyde, likely through their lysine residues. Such intramolecular crosslinks indicate intramolecular folding of the proteins that places two lysine residues or an N-terminus and a lysine residue near each other. Other studies have shown that glutaraldehyde can form intramolecular crosslinks in proteins that leads to altered mobility on SDS-PAGE [16].

t-Darpp oligomerization was analyzed by sedimentation equilibrium [17]. In sedimentation equilibrium, t-Darpp is centrifuged in an analytical ultracentrifuge at a sufficient velocity to move the protein toward the outside of the rotor, yet not high enough to form a pellet. A protein concentration gradient across the centrifuge cell is created by the centrifugal force. At the same time, diffusion acts to oppose the centrifugal force. A concentration equilibrium is reached that can be measured by spectrophotometric absorbance. The concentration distribution detected as a function of distance within the rotor depends only on molecular mass, and is independent of the shape of t-Darpp. Figure 1C shows absorption measurements as a function of distance within the rotor. The data were fitted to a curve that indicates that 96% of the t-Darpp population has a mass of 16 kDa and that the remainder has a mass of 360 kDa. The data largely confirm that the majority of t-Darpp is monomeric and is in agreement with our blue native gel electrophoresis and glutaraldehyde crosslinking studies.

### Shape analysis

Gel filtration chromatography was used to determine the shape of t-Darpp. Two gel filtration columns were used. The Sephadex-50 column effectively separates proteins that are less than 30 kDa. Although recombinant t-Darpp has a calculated mass of 19.6 kDa per monomer, the protein eluted with a mass greater than the column molecular weight cutoff of 30 kDa, assuming a globular shape of the protein (data not shown). This indicates that t-Darpp has an elongated shape.

To explore this further, a Superose 6 gel filtration column (separation range 5–500 kDa) was used to measure the Stokes radius of t-Darpp. The gel filtration chromatograph in Figure 2 shows that t-Darpp eluted between gamma globulin (158 kDa, Stokes radius 5.1 nm) and chicken ovalbumin (44 kDa, Stokes radius 2.8 nm). From these and other mass standards the Stokes radius of t-Darpp was determined to be 4.4 nm. The ratio of the Stokes radius of a protein to the calculated radius of a spherical protein with the same mass can be used to classify the shape of the protein [18]. For t-Darpp, this ratio is 2.4 (Calculation 1S), indicating that t-Darpp is an elongated protein.

**Figure 2.**
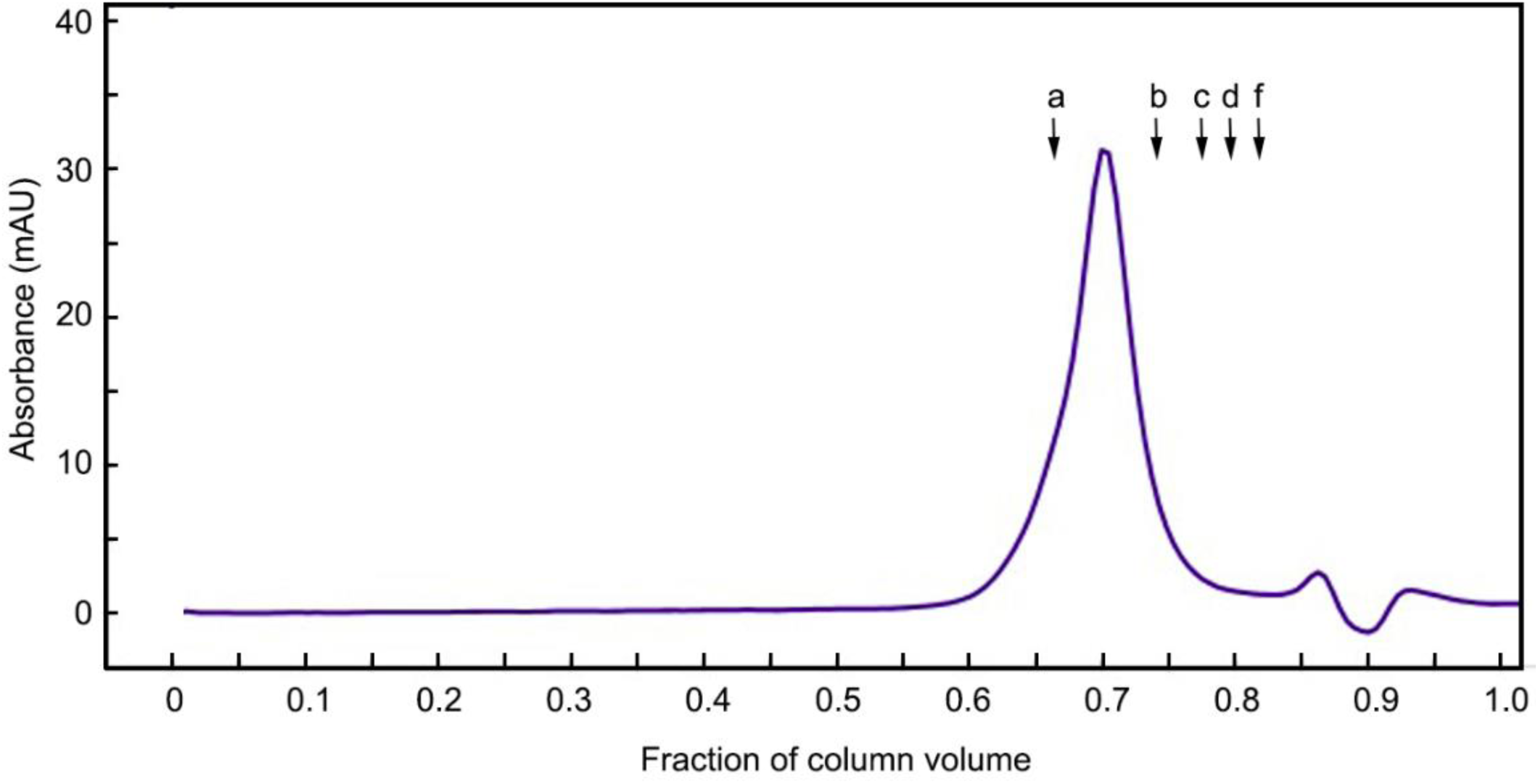
Gel filtration chromatography of t-Darpp. Elution profile of recombinant t-Darpp on Superose 6 column. Elution positions of protein standards indicated as a, 158 kDa (Stokes radius 5.1 nm); b, 44 kDa (Stokes radius 2.8 nm); c, 29 kDa (Stokes radius 2.1 nm); d, 17 kD (Stokes radius 1.9 nm); e, 12 kDa (Stokes radius 1.6 nm).

### Kinases that phosphorylate t-Darpp

A residue critical to t-Darpp activity is T39 (the analogous site within the longer isoform Darpp-32 is T75) because a mutation at this site renders t-Darpp incapable of both, activating the AKT pathway and conferring trastuzumab resistance [1, 2]. CDK1 and CDK5 phosphorylate Darpp-32 at T75 [19]. Figure 3 shows that in *in vitro* kinase reactions, T39 within t-Darpp can be phosphorylated by CDK1 and CDK5. Furthermore, phosphorylation is inhibited by the addition of kinase-specific inhibitors (RO-3306 for CDK1 and Roscovatine for CDK5). Notably, there is a subpopulation of the recombinant t-Darpp that is phosphorylated during its expression in *E. coli*, an observation confirmed by mass-spectrometry (data not shown).

**Figure 3.**
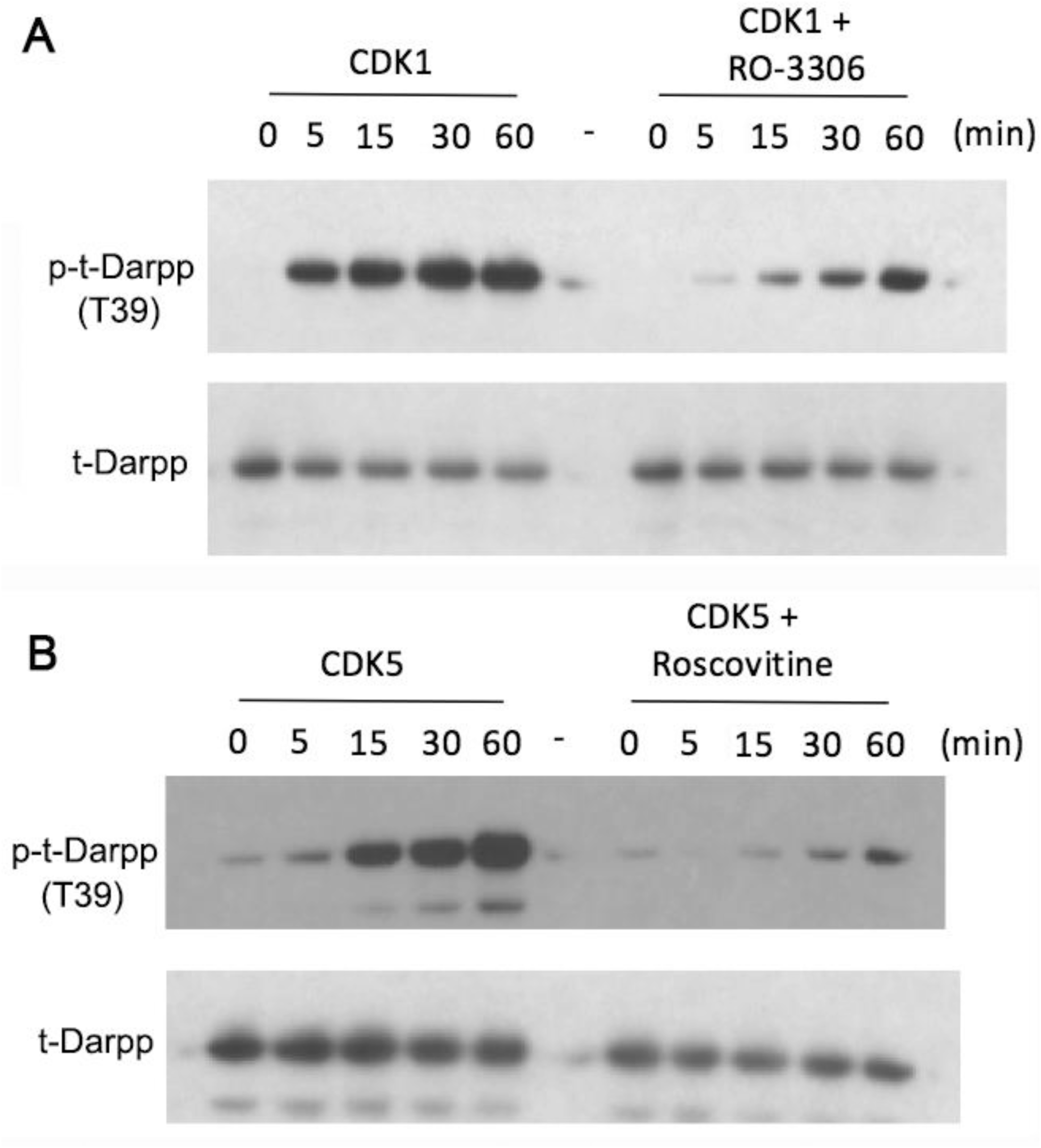
CDK1 and CDK5 phosphorylate t-Darpp *in vitro*. *In vitro* kinase assay performed using recombinant t-Darpp (tDp) and CDK1 (A) or CDK5 (B). Reactions were performed over a period of 1 h with aliquots taken at 0, 5, 15, 30 and 60 min. Inhibitors, RO-3306 and roscovitine were used as controls to inhibit CDK1 and CDK5, respectively. The T39-phosphospecific antibody was used to show phosphorylation at T39 of t-Darpp. H62 was used as a loading control antibody to show the relative level of recombinant t-Darpp in each lane.

### Secondary structure analysis

To measure the secondary structure content of t-Darpp, CD spectroscopy was performed (Figure S1). The CD spectrum was used to determine the percentages of different secondary conformations. Table 1 shows that t-Darpp contains 12% alpha helix, 29% beta strand, 24% beta turn and 35% random coil at ambient temperature. To assess whether the protein structure is sensitive to mild heat, the temperature of the t-Darpp solution was increased from ambient to 50°C for 5 min and allowed to cool slowly back to the ambient temperature. Re-measurement of the heat-treated t-Darpp indicated that the protein secondary structure was not significantly altered: 12% alpha helix, 31% beta strand, 24% beta turn, and 34% random coil.

**Table 1.** Circular dichroism analysis of t-Darpp and Darpp-32

|  | t-Darpp<br>(25°C) | t-Darpp<br>(50°C, 5 min) | Darpp-32<br>(25°C) | Darpp-32<br>(50°C, 5 min) |
| --- | --- | --- | --- | --- |
| alpha helix (%) | 12 | 11 | 18 | 12 |
| beta strand (%) | 29 | 31 | 23 | 32 |
| beta turn (%) | 24 | 24 | 30 | 26 |
| random coil (%) | 35 | 34 | 29 | 29 |

For a comparison, CD analysis was performed on recombinant Darpp-32, which contains an additional 36 residues on the N-terminus (Figure S2). Darpp-32 contains 18% alpha helix, 23% beta strand, 30% beta turn, and 29% random coil (see Table 1). The data indicates that the N-terminal 36 amino acids of Darpp-32 significantly alter the secondary structure composition of the protein by increasing the alpha helical content by 6%, increasing the beta turn content by 6%, decreasing the random coil by 6%, and decreasing the beta strand character by 6%. Unlike t-Darpp, upon brief heat treatment of Darpp-32 there was a significant change in the secondary structure. After heating, Darpp-32 structure was remarkably similar to t-Darpp: 12% alpha helix, 32% beta strand, 26% turn and 29% random coil. The data suggest that the N-terminal region of Darpp-32 is heat labile but the remainder of the protein is not.

### Metal binding analysis

t-Darpp has a calculated isoelectric point of 4.2 and has an acidic segment that stretches from residues 83 to 100 (see Figure 4B). Given these properties, it was investigated whether t-Darpp associates with positively charged metals commonly found in biological systems. Inductively coupled plasma-mass spectrometry (ICP-MS) of recombinant t-Darpp was used to survey a range of metals including Al, Co, Cr, Cu, Fe, Mg, Mg, Ni, Ti, V, and Zn. The detection limit was one metal atom per 10 t-Darpp monomers. Interestingly, none of these metals was detected by ICP-MS. ICP-MS gave a large background of calcium signal. Therefore, graphite furnace atomic adsorption spectroscopy was used to measure the Ca content in t-Darpp. Recombinant t-Darpp was found to contain a significant level of calcium, with an average calcium-to-t-Darpp monomer stoichiometry of 0.57 ± 0.33 (Table 2). To test for alterations in the t-Darpp secondary structure upon metal binding, calcium phosphate and magnesium sulfate were separately spiked into t-Darpp and Darpp-32. CD spectrum analysis (Figures S3, S4) showed no significant change in the secondary structures. Prior incubation with calcium or the calcium chelator EGTA had no effect on t-Darpp mobility on blue native gels (data not shown).

**Figure 4.**
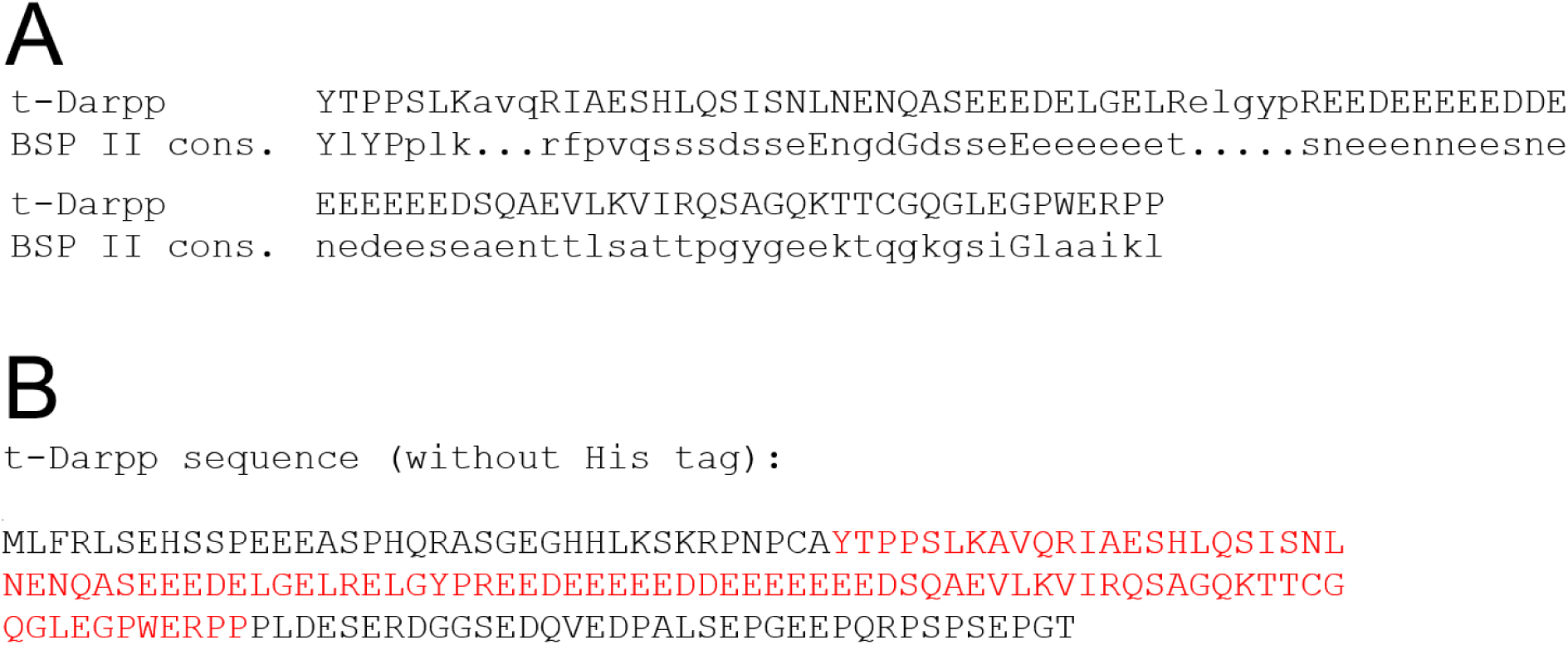
Alignment of t-Darpp with bone sialoprotein II consensus sequence. A. Sequence alignment; B. Entire t-Darpp sequence with residues that align with the bone sialoprotein II consensus sequence presented in underlined font.

**Table 2.** Graphite furnace atomic adsorption spectroscopy analysis

| Trial number | Preparation | Ca/t-Darpp<sup>a</sup> |
| --- | --- | --- |
| 1 | 1 | 0.24 |
| 2 | 1 | 1.00 |
| 3 | 2 | 0.40 |
| 4 | 2 | 0.64 |

<sup>a</sup>Molar ratio

Given that t-Darpp appears to bind to calcium, a motif-searching tool from the GenomeNet suite (National Bioscience Database Center of the Japan Science and Technology Agency) was used to determine potential motifs in t-Darpp (Figure 4). As expected, the output of the motif tool search showed that the highest scoring hit is Darpp-32 (data not shown). The second highest scoring hit is the consensus sequence of bone sialoprotein II (BSP-II) (iE-value = 0.061), a calcium binding protein located in bone and teeth. The region of t-Darpp that aligns with BSP-II is residues 38-131, which contains several Glu and Asp residues, some of which may be key in binding to calcium.

## DISCUSSION

t-Darpp protein is found in normal adult brain striatal cells and in some cancers. t-Darpp activates PKA and increases phosphorylation of AKT. To the best of our knowledge, this is the first *in vitro* study of the t-Darpp protein structure properties. Our results revealed that t-Darpp is monomeric, elongated (Stokes radius 4.4 nm), and binds to calcium ion. t-Darpp has a critical site of phosphorylation at T39 that is required for activation of the AKT pathway and trastuzumab resistance [1, 2]. Our study shows that t-Darpp is phosphorylated by CDK1 and CDK5 at this site. t-Darpp has 12% alpha helix, 29% beta strand, 24% beta turn and 35% random coil character at 25°C and the secondary structure does not appreciably change upon brief mild heat treatment.

Through sedimentation equilibrium analysis, blue native gel electrophoresis, and glutaraldehyde crosslinking we conclude that t-Darpp exists primarily as a monomer and does not form hetero-oligomers with Darpp-32, its longer isoform. Phosphorylation at T39 in t-Darpp is required for its activation of AKT, its effect on trastuzumab resistance, and its stimulation of PKA activity.[1, 7] On the other hand, Darpp-32 is a PKA inhibitor and PP-1 inhibitor, depending on its phosphorylation state [13, 19]. An antagonistic relationship between t-Darpp and Darpp-32 has been reported with regard to both PKA activity and chemoresistance [8]. One possibility to account for these effects is that t-Darpp might alter the Darpp-32 structure, which could promote Darpp-32’s PP-1 inhibitory activity and lead to increased phospho-AKT and/or inhibit Darpp-32’s PKA inhibitory activity and lead to enhanced PKA activity (and possibly promote AKT phosphorylation). The lack of the propensity of t-Darpp to form appreciable levels of hetero-oligomers with Darpp-32 or form homo-oligomers suggests that t-Darpp does not directly alter Darpp-32’s effects on PP-1 or PKA, although it remains possible that *in vivo* oligomer formation could occur and have an antagonistic effect on Darpp-32 function.

The recombinant form of t-Darpp was purified from *E. coli*. Through graphite furnace-atomic adsorption spectroscopy, we show that t-Darpp co-purifies with calcium with a stoichiometry of approximately 0.6 calcium atoms per t-Darpp monomer. The stoichiometry varied somewhat from one t-Darpp preparation to another (Table 2). Calcium-binding properties of proteins can be regulated by phosphorylation [20]. Recombinant t-Darpp prepared from *E. coli* lysates is sub-stoichiometrically phosphorylated as assessed by mass spectrometry (data not shown). It is possible that calcium binding to t-Darpp may depend on its phosphorylation state and that the stoichiometry of calcium to t-Darpp varies depending on the level of t-Darpp phosphorylation in each preparation. Motif analysis of the t-Darpp sequence indicates that its acidic region is similar to the calcium-binding motif of bone sialoprotein II, a mineral nucleating protein produced by osteoblasts and deposited in connective tissue (see Figure 4) [21]. The region of shared similarity between t-Darpp and the bone sialoprotein II consensus sequence includes a large polyglutamic acid segment critical for bone sialoprotein II to bind to calcium [22] and hydroxyapatite (Ca_10_(PO_4_)_6_(OH)_2_) [23, 24]. Other biological metals, including copper, iron, magnesium, manganese, and zinc were not detected in our t-Darpp preparations demonstrating that it is likely that t-Darpp specifically binds to calcium and no other metal.

t-Darpp secondary structure consists of 12% alpha helix, 29% beta strand, 24% beta turn and 35% random coil character at 25°C. After brief heating to 50°C and return to 25°C, the structure does not significantly change. Darpp-32, on the other hand, consists of 18% alpha helix, 23% beta strand, 30% beta turn, and 29% random coil. Brief heat treatment of Darpp-32 and return to 25°C results in a structure similar to t-Darpp: 12% alpha helix, 32% beta strand, 26% beta turn and 29% random coil. NMR structure analysis of Darpp-32 segment 1–118 indicates that this protein has an alpha helix within amino acid residues 22-29 [25]. This alpha helix is transient because it exists only 36% of the time [25] and it is likely to be the region that is heat-labile in our study. This region, required for PP-1 inhibitory activity, is not present in t-Darpp, which is consistent with the absence of a heat-labile structure in t-Darpp.

Our data indicates that t-Darpp is an elongated monomeric protein that is phosphorylated by CDK1 and CDK5. t-Darpp binds to calcium but to no other biological metal. We speculate that calcium binds to the polyglutamic acid region of t-Darpp and that phosphorylation controls its binding to calcium, which may alter its structure. Structure changes may also allow t-Darpp to bind to biological molecules to increase PKA activity and as well as lead to AKT phosphorylation, either through direct interaction with PKA or indirect interaction with one or more regulatory protein(s). t-Darpp does not appreciably form hetero-oligomers with Darpp-32, its longer isoform, so it appears that direct interaction of the two proteins does not account for the antagonistic effects of the two isoforms previously reported. Finding *in vivo* binding partners will shed more light on t-Darpp’s biological activities and give insight into how this protein mediates resistance to trastuzumab.

## EXPERIMENTAL PROCEDURES

### t-Darpp and Darpp-32 subcloning and purification

t-Darpp cDNA (UniProt Accession ID Q9UD71-2) was amplified from t-Darpp/pcDNA3.0/Neo plasmid [8] with primers that include sequences that code for an NcoI restriction site in the upstream primer and an XhoI restriction site in the downstream primer. The downstream primer also included a sequence that codes for a six-His tag at the C-terminus. After treatment with NcoI and XhoI, the cDNA was cloned into NcoI and XhoI sites of the multiple cloning site in the pET-28a plasmid (Novagen). After T4 ligase treatment the plasmid was transformed into competent *E. coli* JM109 cells. DNA sequencing verified that the t-Darpp cDNA shared 100% identity with the wild-type sequence, with the addition of six histidine residues at the C-terminus.

Darpp-32 cDNA (UniProt Accession ID Q9UD71-1) was amplified from Darpp-32/pET-49 plasmid with primers that included sequences that code for a thrombin cleavage site and a six-His tag at the C-terminus. The 5’ end primer contained an NcoI restriction site and the 3’ end primer contained an XhoI restriction site. Amplified Darpp-32 cDNA was cloned into the pET-28a plasmid. After T4 ligase treatment the plasmid was transformed into competent JM109 cells. The sequence added to the C-terminus of Darpp-32 is AAAELALVPRGSLEHHHHHH.

t-Darpp and Darpp-32 expression plasmids were isolated from the JM109 cells and transformed into *E. coli* BL21 DE3 cells. Cells were incubated in Miller’s broth with constant shaking at 37°C until they reached an optical density of 0.6 and treated with 0.5 mM isopropyl β-D-1-thiogalactopyranoside (IPTG) for 16 h at 22°C. Cells were collected by centrifugation at 4000 × g for 20 min at 4°C and supernatant was removed. Cells were frozen at -80°C and lysed at a pellet-to-lysis buffer ratio of 1.7 g to 10 mL. Lysis buffer was 50 mM NaH_2_PO_4_, 500 mM NaCl, 5 mM sodium citrate, and 20 mM imidazole (pH 8.0) or 50 mM HEPES, 500 mM NaCl, and 20 mM imidazole (pH 8.0). Added to the lysis buffer was 6.2 μg/mL RNAse A, 0.5% (v/v) Triton X-100, 1 cOmplete™. ULTRA Mini, EDTA-free, *EASY* pack tablet (Roche), and 1 mM phenylmethylsulfonylfluoride (PMSF). Pelleted cells were resuspended in lysis buffer by vortexing and sonicating with a sonicator (550 Sonic Dismembrator, Fisher Scientific) with six sets of 10 s pulse/10 s cool-down cycles (cool-down temperature was 0°C) at power setting “5.” Lysate was centrifuged twice at 10,000 × g, 4°C for 20 min and supernatant was collected (soluble lysate) after each centrifugation. The soluble lysate was filtered through a 0.22 micron filter.

A 1.5 × 10 cm column containing Ni-NTA resin (Thermo Scientific, 88220) with a bed volume of 0.5 mL was equilibrated with 10 column volumes of water followed by 10 column volumes of wash buffer (20 mM imidazole in buffer A containing 50 mM NaH_2_PO_4_, 500 mM NaCl, 5 mM sodium citrate, 20 mM imidazole, 1 mM PMSF, pH 8.0 or 50 mM HEPES, 500 mM NaCl, 20 mM imidazole, 1 mM PMSF, pH 8.0). Soluble lysate was applied to the column at a flow rate of approximately 1 mL/min. After loading, the column was washed with 10 column volumes of 30 mM imidazole in buffer A. t-Darpp and Darpp-32 were eluted with 40–200 mM imidazole in buffer A. SDS-PAGE analysis indicated that there was no need for further purification for Darpp-32.

To further purify t-Darpp, fractions containing t-Darpp were pooled and mixed 1:1 with buffer B (20 mM L-histidine, 1 mM PMSF, pH 5.5) and loaded onto a 1.5 × 10 cm column packed with Q-Sepharose (Fastflow, Pharmacia, 17-0510-01) containing a bed volume of 0.5 mL. The column was washed with 0.2 M NaCl in buffer B and t-Darpp was eluted with 0.4 M NaCl in buffer B. t-Darpp and Darpp-32 were concentrated with an Amicon Ultra 2-mL centrifugal filter (EMD Millipore; 10 MW cutoff) and buffer was exchanged to 50 mM HEPES, 150 mM NaCl, 1 mM EDTA, 5% glycerol, 1 mM DTT (pH 7.0) and stored at -20°C. Relative protein band densities in SDS-PAGE gels were measured with ImageJ software (Version 1.51a) [26].

### Blue native gel electrophoresis

Blue native gel electrophoresis was performed as described [27]. Briefly, purified t-Darpp was mixed with an equal volume of native gel loading buffer (50 mM Bis-Tris/HCl, pH 7.0, 15% (v/v) glycerol) and electrophoresed using a cathode buffer (15 mM Bis-Tris/HCl, pH 7.0, 50 mM Tricine, 0.002% (w/v) Coomassie Blue) and an anode buffer (50 mM Bis-Tris/HCl, pH 7.0). To avoid overheating, the gel was run at 100 V at 4°C. Samples were analyzed on discontinuous gel consisting of 5% stacking gel and 8% resolving gel prepared using a 3x-Gel Buffer (150 mM Bis-Tris/HCl, pH 7.0). Native molecular weight markers (Invitrogen™ Novex™ NativeMark™, Thermo Fisher Scientific) and protein samples were detected by accumulation of Coomassie blue dye within the proteins during electrophoresis.

### CD analysis

CD analysis was performed at room temperature on a JASCO J-815 CD spectrometer (JASCO Inc., Tokyo, Japan) in a 1-mm path quartz cell. Spectra were collected between 260 and 190 nm at 0.1 nm intervals. The bandwidth was 1 nm and the scan rate was 100 nm/min. Samples were dissolved in 50 mM sodium phosphate, pH 7.4. Temperature was controlled via a Fisher Scientific ISOTEMP 102 water bath (Thermo Fisher Scientific, Waltham, USA). Eight scans were collected per sample and averaged. Protein samples were diluted so that High Tension (HT) voltage traces were less than 500 V. Final concentration of t-Darpp and Darpp-32 was 0.12 mg/mL. Data were formatted and analyzed by CDPro software (http://sites.bmb.colostate.edu/sreeram/CDPro/) [28]. Experimental data nearly matched reconstructed data created by the CDSSTR and CONTIN/LL programs. Percentages of proteins classified as alpha helix, beta strand, beta turn and random coil were calculated by averaging the outputs from these two software programs. For metal spiking experiments, calcium was added into the protein solution in the form of Ca(H_2_PO_4_)_2_ and magnesium was added as MgSO_4_.

### Glutaraldehyde crosslinking

Glutaraldehyde (Fisher Scientific) was added to a final concentration 0.1% (v/v) and 1% (v/v) to protein solutions and incubated 30 min at 25°C. Samples were quenched by the addition of sample buffer (0.125 M Tris, pH 6.8, 20% (v/v) glycerol, 0.02% (w/v) bromphenol blue, 20% (w/v) sodium docecyl sulfate, 1.4 M 2-mercaptoethanol, pH 6.8) and separated on a 10% cross-linked polyacrylamide gel [29].

### In vitro kinase assay

Roscovitine (sc-24002) and RO-3306 (sc-358700) were purchased from Santa Cruz Biotechnology (Dallas, TX). Recombinant t-Darpp was used as a substrate for CDK1 and CDK5 phosphorylation. CDK1 (C22-18G) and CDK5 (C33-10G-05) were purchased from Signal Chem Pharmaceuticals (Richmond BC, Canada). CDK1 or CDK5 was diluted to a final concentration of 5 μg/mL in kinase dilution buffer (K23-09, Signal Chem Pharmaceuticals). The reactions were pre-incubated with t-Darpp for 10 min at 30°C. The kinase reaction was initiated by the addition of 200 μM ATP (final concentration) and continued at 30°C over a period of 60 min. Reactions were quenched by the addition of sample buffer and boiled for 5 min. RO-3306 or roscovitine at 10 μM was used to inhibit CDK1 or CDK5 activity, respectively. These inhibitors were added directly to the kinase reaction during the preincubation period. Samples were analyzed by Western analysis as described below.

### Western blot analysis

Proteins were separated by 10% SDS-polyacrylamide gel electrophoresis and transferred onto a nitrocellulose membrane. Blocking as well as incubation with primary and secondary antibodies were performed in 5% (w/v) non-fat dry milk. Primary antibody against phospho-Darpp-32/t-Darpp (T39) (#2301) was from Cell Signaling. H62 (sc-11365), an antibody that recognizes both Darpp-32 and t-Darpp, was from Santa Cruz Biotechnology. Horseradish peroxidase-conjugated anti-rabbit IgG secondary antibody was purchased from Cell Signaling. Chemiluminscence was produced using the Amersham Plus kit and captured on X-ray film.

### Gel filtration analysis

A G-50 Sephadex column (2 × 15 cm) was used initially to estimate the Stokes radius of t-Darpp. Standards used were bovine aprotinin (6.5 kDa and 1.48 nm Stokes radius), bovine cytochrome c (12.4 kDa and 1.6 nm Stokes radius), and bovine carbonic anhydrase (29 kDa and 2.1 nm Stokes radius) [30, 31]. The column was run at a flow rate of 1 mL/min in 150 mM HEPES (pH 7.0) containing 1 mM dithiothreitol (DTT). A second column was used to measure the Stokes radius. Superose 6 column (Pharmacia Biotech 6 HR 10/30) was equilibrated with 50 mM HEPES (pH 7.0), 150 mM NaCl, 1 mM EDTA, 1 mM dithiothreitol, and 5% glycerol at a flow rate of 0.3 mL/ min. In addition to the standards listed above, the following standards were used: thyroglobulin (670 kDa and 8.6 nm Stokes radius), bovine gamma globulin (158 kDa and 5.1 nm Stokes radius), chicken ovalbumin (44 kDa and 2.8 nm Stokes radius), and horse myoglobin (17 kDa and 1.9 nm Stokes radius). A 50 μL sample of t-Darpp (4 mg/mL) was applied to the column. Classification of t-Darpp as a highly elongated protein was based on calculated mass and experimentally derived Stokes radius using the method of Erikson [18].

### Graphite furnace–atomic adsorption spectrometry

Calcium analyses were performed on an atomic absorption spectrophotometer with a graphite furnace (GFAA-7000, Shimatzu). An autosampler (ASC-7000, Shimatzu) was used for ensuring good precision. An absorption detection wavelength of 422.7 nm was used and signal was corrected by the BGC-D2 mode. A seven-stage furnace treatment program was used (Table S1). For each sample, a minimum of four injections were performed to achieve a relative standard deviation of 5%. A calibration curve was used for the determination of calcium concentration in the diluted protein solution.

Standard calcium solutions were prepared by diluting from a single standard calcium solution (1000 ppm in 2% HNO_3_, Perkin Elmer) with 0.1% metal-free HNO_3_ (GFS Chemicals Inc.; Powell, OH) to desired concentrations. Trace metal-free vials were used for sample dilutions. The protein sample solutions were also diluted with 0.1% HNO_3_ prepared from trace metal grade concentrated HNO3 (67-70%, Fisher Scientific) and deionized water of 18.2 MΩ·cm (Milli-Q system, Millipore).

### Inductively coupled plasma-mass spectrometry (ICP-MS)

All samples for ICP-MS analysis were acidified with Optima grade nitric acid and analyzed at IIRMES, California State University Long Beach. Triplicate blank samples were prepared in the same manner as all samples for analysis and the average concentration of the various metals in the blanks was subtracted from all samples.

### Sedimentation equilibrium analysis

Sedimentation equilibrium runs were conducted on a Beckman Optima XL-A analytical ultracentrifuge. Protein sedimentation was monitored at 280 nm at 4°C. t-Darpp at 0.58 μg/μL in phosphate buffered saline was centrifuged in a 12-mm path length six-sector cell. Sedimentation equilibrium profile was measured at 9,000 rpm. Data were fitted to a single exponential curve using Beckman global analysis software to determine the weighted average of protein masses.

## ACKNOWLEDGMENTS

This research was supported by a grant from the National Institute of General Medical Science 1RO1GM105898. FZ also acknowledges support from the NSF-Center for Research Excellence in Science and Technology Program (NSF HRD 1547723) and XW thanks partial support from the National Key Basic Research Program (2014CB744502). The authors are indebted to Drs. Rich Gossett, Yuan Yu Lee, and Alexander Long at the IIRMES laboratory at California State University Long Beach for ICP-MS analyses of t-Darpp metal content.

## AUTHOR CONTRIBUTIONS

JM designed the study and wrote the paper. FZ and SEK also contributed to writing the manuscript. PM, EP, AC, and YF purified recombinant t-Darpp and YF subcloned and purified Darpp-32. PM performed blue native gel electrophoresis and Stokes radius measurments, MLP conducted sedimentation equilibrium centrifugation studies, DJ and EM conducted graphite furnace-atomic adsorption spectroscopy analysis, XW and YF conducted CD experiments, YF conducted cross-linking experiments, and ED performed t-Darpp phosphorylation experiments. All authors reviewed the results and approved the final version of the manuscript.

## REFERENCES

1. Vangamudi, B., Peng, D. F., Cai, Q., El-Rifai, W., Zheng, W. & Belkhiri, A. (2010) t-DARPP regulates phosphatidylinositol-3-kinase-dependent cell growth in breast cancer, Molecular cancer. 9, 240.

2. Hamel, S., Bouchard, A., Ferrario, C., Hassan, S., Aguilar-Mahecha, A., Buchanan, M., Quenneville, L., Miller, W. & Basik, M. (2010) Both t-Darpp and DARPP-32 can cause resistance to trastuzumab in breast cancer cells and are frequently expressed in primary breast cancers, Breast cancer research and treatment. 120, 47-57.

3. El-Rifai, W., Smith, M. F., Jr., Li, G., Beckler, A., Carl, V. S., Montgomery, E., Knuutila, S., Moskaluk, C. A., Frierson, H. F., Jr. & Powell, S. M. (2002) Gastric cancers overexpress DARPP-32 and a novel isoform, t-DARPP, Cancer Res. 62, 4061-4.

4. Beckler, A., Moskaluk, C. A., Zaika, A., Hampton, G. M., Powell, S. M., Frierson, H. F., Jr. & El-Rifai, W. (2003) Overexpression of the 32-kilodalton dopamine and cyclic adenosine 3′,5′-monophosphate-regulated phosphoprotein in common adenocarcinomas, Cancer. 98, 1547-51.

5. Straccia, M., Carrere, J., Rosser, A. E. & Canals, J. M. (2016) Human t-DARPP is induced during striatal development, Neuroscience. 333, 320-30.

6. Belkhiri, A., Dar, A. A., Peng, D. F., Razvi, M. H., Rinehart, C., Arteaga, C. L. & El-Rifai, W. (2008) Expression of t-DARPP mediates trastuzumab resistance in breast cancer cells, Clin Cancer Res. 14, 4564-71.

7. Gu, L., Lau, S. K., Loera, S., Somlo, G. & Kane, S. E. (2009) Protein kinase A activation confers resistance to trastuzumab in human breast cancer cell lines, Clin Cancer Res. 15, 7196-206.

8. Gu, L., Waliany, S. & Kane, S. E. (2009) Darpp-32 and its truncated variant t-Darpp have antagonistic effects on breast cancer cell growth and herceptin resistance, PloS one. 4, e6220.

9. Williams, K. R., Hemmings, H. C., Jr., LoPresti, M. B., Konigsberg, W. H. & Greengard, P. (1986) DARPP-32, a dopamine- and cyclic AMP-regulated neuronal phosphoprotein. Primary structure and homology with protein phosphatase inhibitor-1, J Biol Chem. 261, 1890-903.

10. Berger, B., Febvret, A., Greengard, P. & Goldman-Rakic, P. S. (1990) DARPP-32, a phosphoprotein enriched in dopaminoceptive neurons bearing dopamine D1 receptors: distribution in the cerebral cortex of the newborn and adult rhesus monkey, J Comp Neurol. 299, 327-48.

11. Ouimet, C. C., Miller, P. E., Hemmings, H. C., Jr., Walaas, S. I. & Greengard, P. (1984) DARPP-32, a dopamine- and adenosine 3′: 5′-monophosphate-regulated phosphoprotein enriched in dopamine-innervated brain regions. III. Immunocytochemical localization, J Neurosci. 4, 111-24.

12. Hemmings, H. C., Jr., Greengard, P., Tung, H. Y. & Cohen, P. (1984) DARPP-32, a dopamine-regulated neuronal phosphoprotein, is a potent inhibitor of protein phosphatase-1, Nature. 310, 503-5.

13. Le Novere, N., Li, L. & Girault, J. A. (2008) DARPP-32: molecular integration of phosphorylation potential, Cell Mol Life Sci. 65, 2125-7.

14. Yun, S. M., Yoon, K., Lee, S., Kim, E., Kong, S. H., Choe, J., Kang, J. M., Han, T. S., Kim, P., Choi, Y., Jho, S., Yoo, H., Bhak, J., Yang, H. K. & Kim, S. J. (2014) PPP1R1B-STARD3 chimeric fusion transcript in human gastric cancer promotes tumorigenesis through activation of PI3K/AKT signaling, Oncogene. 33, 5341-7.

15. Shukolyukov, S. A. (2009) Aggregation of frog rhodopsin to oligomers and their dissociation to monomer: application of BN- and SDS-PAGE, Biochemistry (Mosc). 74, 599-604.

16. Ross, D. C. & McIntosh, D. B. (1987) Intramolecular cross-linking of domains at the active site links A1 and B subfragments of the Ca2+-ATPase of sarcoplasmic reticulum, J Biol Chem. 262, 2042-9.

17. Bothwell, M. A., Howlett, G. J. & Schachman, H. K. (1978) A sedimentation equilibrium method for determining molecular weights of proteins with a tabletop high speed air turbine centrifuge, J Biol Chem. 253, 2073-7.

18. Erickson, H. P. (2009) Size and shape of protein molecules at the nanometer level determined by sedimentation, gel filtration, and electron microscopy, Biol Proced Online. 11, 32-51.

19. Bibb, J. A., Snyder, G. L., Nishi, A., Yan, Z., Meijer, L., Fienberg, A. A., Tsai, L. H., Kwon, Y. T., Girault, J. A., Czernik, A. J., Huganir, R. L., Hemmings, H. C., Jr., Nairn, A. C. & Greengard, P. (1999) Phosphorylation of DARPP-32 by Cdk5 modulates dopamine signalling in neurons, Nature. 402, 669-71.

20. Alsheikh, M. K., Svensson, J. T. & Randall, S. K. (2005) Phosphorylation regulated ion-binding is a property shared by the acidic subclass dehydrins, Plant, Cell & Environment. 28, 1114-1122.

21. Ganss, B., Kim, R. H. & Sodek, J. (1999) Bone sialoprotein, Crit Rev Oral Biol Med. 10, 79-98.

22. Chen, Y., Bal, B. S. & Gorski, J. P. (1992) Calcium and collagen binding properties of osteopontin, bone sialoprotein, and bone acidic glycoprotein-75 from bone, J Biol Chem. 267, 24871-8.

23. Stubbs, J. T., 3rd, Mintz, K. P., Eanes, E. D., Torchia, D. A. & Fisher, L. W. (1997) Characterization of native and recombinant bone sialoprotein: delineation of the mineral-binding and cell adhesion domains and structural analysis of the RGD domain, J Bone Miner Res. 12, 1210-22.

24. Tye, C. E., Rattray, K. R., Warner, K. J., Gordon, J. A., Sodek, J., Hunter, G. K. & Goldberg, H. A. (2003) Delineation of the hydroxyapatite-nucleating domains of bone sialoprotein, J Biol Chem. 278, 7949-55.

25. Dancheck, B., Nairn, A. C. & Peti, W. (2008) Detailed structural characterization of unbound protein phosphatase 1 inhibitors, Biochemistry. 47, 12346-56.

26. Schneider, C. A., Rasband, W. S. & Eliceiri, K. W. (2012) NIH Image to ImageJ: 25 years of image analysis, Nat Methods. 9, 671-5.

27. Schagger, H., Cramer, W. A. & von Jagow, G. (1994) Analysis of molecular masses and oligomeric states of protein complexes by blue native electrophoresis and isolation of membrane protein complexes by two-dimensional native electrophoresis, Analytical biochemistry. 217, 220-30.

28. Sreerama, N. & Woody, R. W. (2000) Estimation of protein secondary structure from circular dichroism spectra: comparison of CONTIN, SELCON, and CDSSTR methods with an expanded reference set, Analytical biochemistry. 287, 252-60.

29. Laemmli, U. K. (1970) Cleavage of structural proteins during assembly of the head of the bacteriophage T4., Nature. 277, 680-685.

30. Mohr, D., Frey, S., Fischer, T., Guttler, T. & Gorlich, D. (2009) Characterisation of the passive permeability barrier of nuclear pore complexes, EMBO J. 28, 2541-53.

31. Potschka, M. (1987) Universal calibration of gel permeation chromatography and determination of molecular shape in solution, Analytical biochemistry. 162, 47-64.

